# Effects of Natural Lithium and Lithium Isotopes on Voltage Gated Sodium Channel Activity in SH-SY5Y and IPSC Derived Cortical Neurons

**DOI:** 10.1101/2025.05.28.656602

**Authors:** Irina Bukhteeva, James D. Livingstone, Kartar Singh, Evgeny V. Pavlov, Michael A. Beazely, Michel J. P. Gingras, Zoya Leonenko

## Abstract

Although lithium (Li) is a widely used treatment for bipolar disorder, its exact mechanisms of action remain elusive. Research has shown that the two stable Li isotopes, which differ in their mass and nuclear spin, can induce distinct effects in both in vivo and in vitro studies. Since sodium (Na^+^) channels are the primary pathway for Li^+^ entry into cells, we examined how Li^+^ affects the current of Na^+^ channels using whole-cell patch-clamp techniques on SH-SY5Y neuroblastoma cells and human iPSC-derived cortical neurons. Our findings indicate that mammalian Na^+^ channels in both neuronal models studied here display no selectivity between Na^+^ and Li^+^, unlike previously reported bacterial Na^+^ channels. We observed differences between the two neuronal models in three measured parameters (*V*_half_, *G*_max_,*z*). We saw no statistically significant differences between any ions in SHSY-5Y cells, but small differences in the half-maximum activation potential (*V*_half_) between Na^+^ and ^6^Li^+^ and between ^7^Li^+^ and ^6^Li^+^ were found in iPSC-derived cortical neurons. Although Na^+^ channels are widely expressed and important in neuronal function, the very small differences observed in this work suggest that Li^+^ regulation through Na^+^ channels is likely not the primary mechanism underlying Li^+^ isotope differentiation.

## Introduction

Lithium (Li), administered in the form of lithium carbonate or citrate salts, has been a forefront medication in the treatment of bipolar disorder for decades^1^. Despite its long-standing clinical use and the systematic exploration of many different targets and pathways^2^, a full understanding of all the mechanisms of Li action is still lacking.

Lithium has two stable isotopes, ^6^Li and ^7^Li, which differ in their atomic mass and nuclear spin. Specifically, ^6^Li has an atomic mass of 6.0151223 atomic mass units (amu) and a nuclear spin of 1, while ^7^Li has an atomic mass of 7.016004 amu and a nuclear spin of 3/2. There is recent interest in the possibility of distinct effects of the two Li isotopes in vitro and in cellular, and animal-based assays. Intriguingly, some studies have shown a large Li isotope effect, including in animal behaviour^3^, neuronal electrical responses^4^ and mitochondrial membrane permeability^5^. On the other hand, others have shown smaller differences^6–11^ or no differences between the isotopes^5,10,12,13^ in different systems. These findings leave unsettled the question of whether distinct Li isotope effects do indeed exist and, if so, raise questions regarding the molecular targets and the mechanisms that may be responsible for the Li^+^ isotope effects that have been reported.

One of the primary pathways for the entry of the lithium ion (Li^+^) into cells is through sodium (Na^+^) channels^14–18^. All previous studies exploring how Li^+^ affects Na^+^ channels have been carried out with natural Li^+^, which consists of approximately 92% of the ^7^Li^+^ isotope, with ^6^Li^+^ making up the other 8% of the natural isotopic abundance of lithium. Given the recognized ability of Li^+^ to enter cells through Na^+^ channels, it is natural to ask how the two Li^+^ isotopes interact with these channels. Here, we focus on two key questions: (1) How do Na^+^ and natural Li^+^ compete, interact, or interfere when passing through Na^+^ channels? (2) Are there any differences between the passage of the two Li isotopes through Na^+^ channels that could help explain observed downstream biological effects?^3–5^

Na^+^ channels are found in eukaryotic and prokaryotic cells. Eukaryotic Na^+^ channels are large, consisting of complex proteins composed of four homologous domains connected by long intracellular linkers whereas prokaryotic Na^+^ channels are formed by the co-assembly of four identical subunits^19,20^. Bacterial Na^+^ channels, with their solved crystal structures in either open or closed state, offer insights into ion selectivity, gating, and drug interaction mechanisms, making them useful models for the more complex eukaryotic counterparts^19^. Some studies on bacterial Na^+^ channels have revealed that the substitution of Na^+^ by natural Li^+^ leads to a decrease of ionic current in these channels^17,18^. On the other hand, a study of rat skeletal muscle has shown that there is no difference between Na^+^ and Li^+^ ionic currents^15^. Given this puzzling difference in terms of Li^+^ interactions with bacterial and mammalian channels^21^, we opted to study mammalian Na^+^ channels since they appear to have been rather incompletely investigated in terms of their behavior under natural (isotopic abundance) Li^+^ exposure and which, to the best of our knowledge, have never been studied in terms of their response under the two different Li^+^ isotopes.

We chose two model cell lines – SH-SY5Y human neuroblastoma cells and human induced pluripotent stem cell (iPSC)-derived neurons. In vitro models, such as the well-studied and characterized SH-SY5Y cell line, facilitate reproducible studies and help explore the pathophysiological mechanisms of neurodegenerative diseases^22^. In addition, iPSC-derived cortical neurons hold promise for bridging the gap between human studies and model organisms, aiding in the understanding of mechanisms and identifying therapeutic strategies for neurological and psychiatric diseases^23^ and offering significant potential for advancing neuroscience research^24,25^.

As discussed above, since Na^+^ channels are an obvious first upstream molecular target of Li^+^ ions as these enter the cell, we investigated how the two Li^+^ isotopes pass through Na^+^ channels in SH-SY5Y human neuroblastoma cells and iPSC-derived neurons using the whole-cell patch-clamp electrophysiology method. We investigated two questions: First, we assessed whether mammalian Na^+^ channels support smaller currents in the presence of Li^+^ ions as was found for bacterial ones^17,18^. Second, we tested whether the presence of Li^+^ isotopes changed various aspects of Na^+^ channel activity and, consequently, neuronal cell excitability, in a way that could help rationalize the large difference between Li^+^ isotope effects reported in some animal behaviour and tissue studies^3,4^.

## Results

### Experimental setup, measurement procedure and analysis

All cells were voltage □ clamped in the whole □ cell configuration. Current-voltage data were obtained by applying a series of 16 consecutive voltage steps, from a holding potential of − 100 mV to +50 mV in +10 mV increments generating 16 corresponding current traces [see lower part of Figure **1c)**]. Each protocol run, including buffer switching, lasted approximately one minute. As the membrane potential becomes progressively more depolarized, more and more Na^+^ channels open, increasing the measured current. We found that the majority of Na^+^ channels in the cell opened at approximately -40 mV in iPSC-derived neurons and at about -20 mV in SH-SY5Y cells. Data were always recorded during continuous perfusion of the clamped cell with the extracellular solution, using a gravity-driven arrangement and at the rate of ∼1 ml/min [Figure **1a)**].

**Fig. 1:**
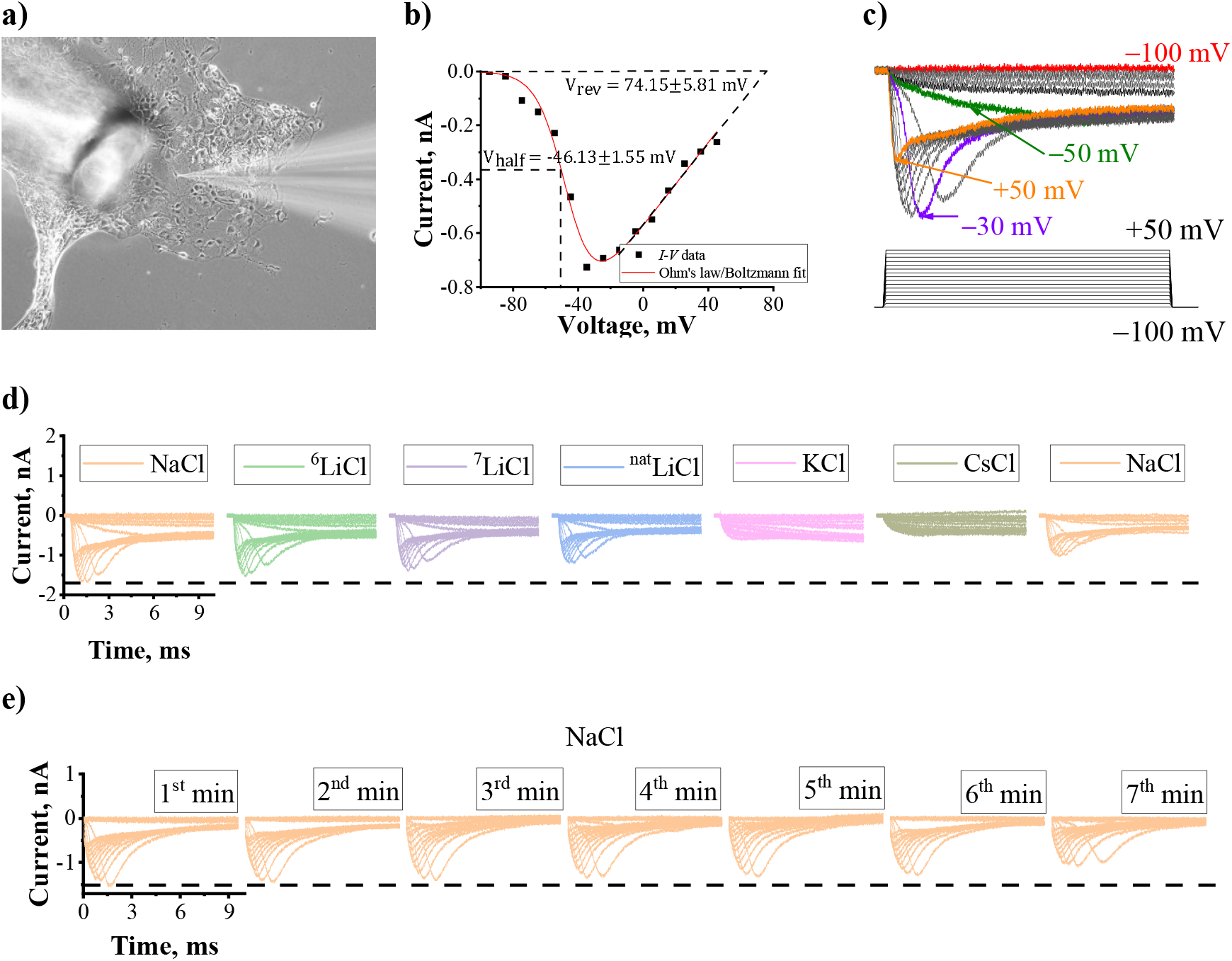
Figure 1. Experimental setup and data analysis. **a)** Experiment setup – a phase contrast image of a patch pipette (on the right) attached to the membrane of an iPSC-derived cortical neuron with the single-cell perfusion tube for extracellular buffer exchange on the left. **b)** Example of *I*–*V* data for one experiment (*n* = 1) analyzed with (Eq. 1): and are shown on the Figure, *G*_max_ = 0.01 0.00 *μ* S, *z* = 0.22 0.03. **c)** Top: Representative traces with colored currents at selected voltage steps to show signal evolution. Bottom: Stimulating protocol used: consecutive sixteen voltage steps from -100 mV to +50 mV with +10 mV increments. **d)** Representative traces of current recordings for one sequence of timeline-ordered perfusions, from left to right: NaCl; ^6^LiCl; ^7^LiCl; ^nat^LiCl; KCl; CsCl; NaCl. For all ionic solutions shown, the time-dependence of the current traces resulting from the consecutive sixteen +10 mV voltage steps is shown. The dashed line indicates the maximum current achieved during the first NaCl perfusion. **e)** Representative current traces for recording in NaCl buffer over 7 minutes to represent rundown in the same buffer. The dashed line indicates the maximum current achieved during the first NaCl perfusion. All examples in this figure are shown for iPSC-derived cortical neurons.

The permeability to different cations was tested by recording an *I*–*V* sequence, first in a control Na^+^ solution, and then, during extracellular perfusion with a sequence of various cations solutions and, finally, again with a control Na^+^ solution (see Figure **1d)** for the sequence of cation solutions used). Henceforth, the notation ^6^LiCl, ^7^LiCl and ^nat^LiCl refers to the various Li chloride salts considered with one of the ^6^Li, ^7^Li or natural isotopic abundance, respectively (see Materials and Methods section).

A current was observed during cell perfusion with 135 mM NaCl (peach), 135 mM ^6^LiCl (green), 135 mM ^7^LiCl (violet), and 135 mM ^nat^LiCl (blue) but was not observed during perfusion with 135 mM KCl (pink) or 135 mM CsCl (bronze). Since the isotope effect on Na^+^ channels was not studied before, we decided to perform perfusions with ^6^LiCl and ^7^LiCl in varying sequences from replicate to replicate to rule out possible order-dependent effects. Perfusion with NaCl was always performed first and last (to check the patch stability and account for the so-called rundown (i.e. the decrease of current amplitude over time), ^nat^LiCl was kept in the fourth place, while KCl and CsCl, which were used as controls, were perfused at the very end of the cation sequence before the terminating NaCl perfusion experiment [see Figure **1d)**].

We anticipated that, if they were to occur, differences might manifest in one of two ways: either as distinct and prominent effects (similar to those observed in behavioral^3^ or fEPSP studies^4^) or as subtle differences (potentially explained by the isotopic mass difference). In the former case, any reasonable experimental design should be sufficient to capture these differences. However, for subtle differences, it is crucial to observe the Li^+^ isotope effects within the same system to ensure reliability. This is why we followed a well-established and employed procedure^15,17,18^, with a sequence of ion perfusions within a single cell to rule out any possible artifacts induced by cell-to-cell differences. On top of this, we decided to add “order randomization” for ^6^LiCl and ^7^LiCl to account for any possibility of order dependence. Upon comparing two different orders of Li^+^ isotopes, we found no differences between the groups (see Supplementary Materials, Figure S2), so further we report all the replicates together.

The peak currents generally decreased progressively over the run time of the experiment for a given patched cell (a phenomenon referred to as “rundown”). As shown in Figure **1d)**, the leftmost and the rightmost subpanels (peach traces) display a reduction in maximum current amplitude, although buffer conditions are the same. For example, here, the maximum current on the leftmost (NaCl) subpanel of Figure **1?)** is about -1.75 nA (around *t*=1.5 ms) while the maximum current on the rightmost subpanel of Figure **1c)** is approximately -1 nA (around *t*=1.5 ms), thus amounting to a reduction of 0.75 nA in current amplitude. To verify that such rundown happens due to the overall duration of the recording period and perfusion stress, and is not due to different cations (that could potentially disturb, thereafter and through their perfusion, the functionality of the channels), we performed a baseline experiment using perfused voltage-clamped cell with solely NaCl without changing to other cations [see Figure **1e**)]. As stated above, about one minute is required to complete and record one protocol consisting of 16 voltage steps from -100 mV to +50 mV. The leftmost trace in Figure **1e)** corresponds to the first minute window (*t*=0 – 1 min) recording of a patched cell with a maximum peak current of about -1.5 nA (around *t*=1.5 ms), the rightmost trace corresponds to the seventh minute window (*t*=6–7 min) of recording on the same cell, displaying a maximum peak current of approximately -1.0 nA (around *t*=1.5 ms), thus indicating a decrease of 0.5 nA in peak current over the 7 minutes with solely NaCl being perfused. It is worth mentioning that the rundown was found to be different for different cells (see examples in Supplementary Material, Figure S1). Therefore, the final signal normalization was done using the first and last recordings in NaCl buffer for a given measured patched cell, rather than normalizing to an averaged rundown across cells. The rundown was interpolated using the first and last recordings with the NaCl buffer and all currents in other buffers were adjusted (i.e. rescaled) according to this interpolation. After data collection and rundown collection, fitting the decay of current to a linear equation^18^ (see Material and Methods section. Electrophysiology subsection for details), the current amplitude was extracted for each I vs time trace. The *I* − *V* characteristics were then plotted for each cell and fitted with (Eq. 1) as illustrated in Figure 1b) for one cell subject to one of the salts considered. From the *I* − *V* characteristic for a given cell, four characteristics were extracted from these fittings), that is *V*_rev_, *V*_half_, *G*_max_ and *z* for each cell. These quantities were then averaged over n replicates and used for further comparison. It is important to note that *V*_rev_ was not experimentally observed but extrapolated based on Ohm’s law. This is further discussed in Materials and Methods and the extrapolated values are reported in the Supplementary Materials. Figure S3.

### Na^+^ and Li^+^ permeability of Na^+^ channels in SH-SY5Y cells

To plot the *I* − *V* channel profiles for Na^+^ and Li^+^ ions, the cunent magnitudes were normalized to the maximum inward current measured in the Na^+^ condition [Figure 2a)]. To obtain a better representation of *V*_half_, the *G* − *V* profiles were also calculated and plotted with normalization to maximum conductance in Na^+^ buffer [Figure 2b)]. To compare these profiles between each other and to account for the unique Na^+^ channel amount in each cell as well as the uniqueness of the patch, characteristics were plotted for each cell and fitted with (Eq. 1) [Figure 1b)] to extract channel parameters - *F*_half_, *G*_max_,*z*. As can be seen from Figure 2. we found no difference in average *I* − *V* or *G* − *V* channel profiles or in any of the channel parameters reported in panels c) to e).

**Fig. 2:**
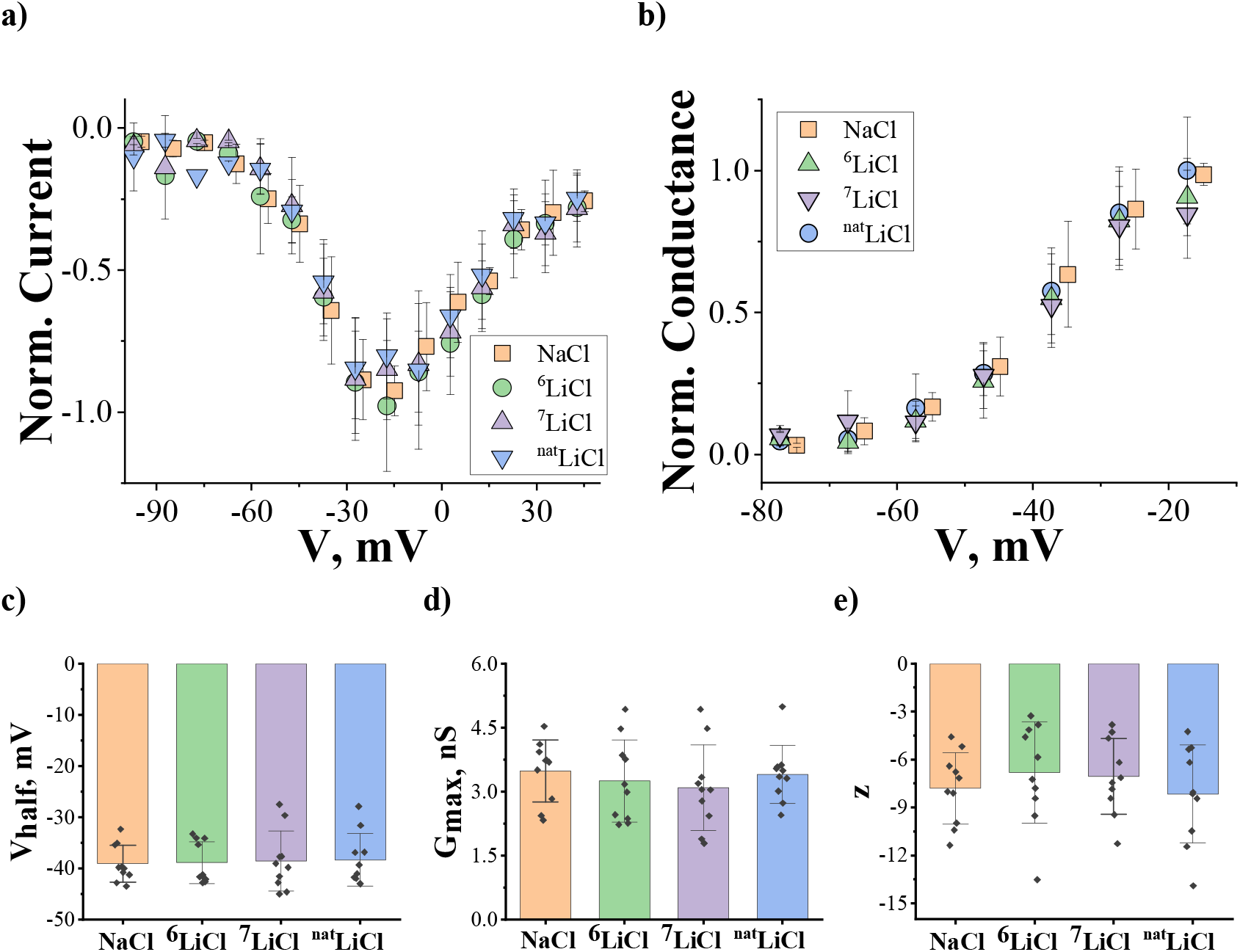
Normalized current/conductance profiles and Na^+^ channel parameters from SH-SY5Y cells upon constant perfusion of NaCl, ^6^LiCl, ^7^LiCl, ^nat^LiCl. **Figure 2. a)** Corresponding *I*−*V* profiles for Na and Li ions, where cunent magnitudes were nonnalized to the maximum inward cunent measured solely under Na^+^ perfusion, **b)** Conesponding *G* − *V* profiles for the Na^+^ and Li” ions, where conductance magnitudes were nonnalized to the maximum conductance measured solely under Na^+^ perfusion, **c)** *V*_half_ − the half-maximum activation potential, **d)** *G*_max_ − the maximal conductance, **e)** *z* − the apparent valence of the gating charge. The symbols in panels a) and b) and columns in panels **c)-e)** represent the mean, the data point (black dots) in panels **c)-e)** represent individual measurements, and bars represent standard deviation (SD). Statistical significance for individual cells was evaluated using one-way repeated measures ANOVA. The number of replicates (individual cells successfully patched) is *n* = 10.

### Na^+^ and Li^+^ permeability of Na^+^ channels in iPSC-derived cortical neurons

Following the SH-SY5Y cells study which we used as a model for comparison, we repeated our experiments on iPSC-derived cortical neurons to investigate how Na^+^ channels in those cells respond to Li ions. We found a significant difference in *V*_half_ between Na^+^ and ^6^LiCl groups and between ^6^LiCl and ^nat^LiCl groups (with a significance level of *p* < 0.05) as shown in Figure 3c). As presented in Figure 3, we found no difference in average *I* — *V* or *G* — *V* channel profiles [panels **a)** and **b)**] as well as in *G*_max_ and *z* [panels **d)** and **e)**].

**Fig. 3:**
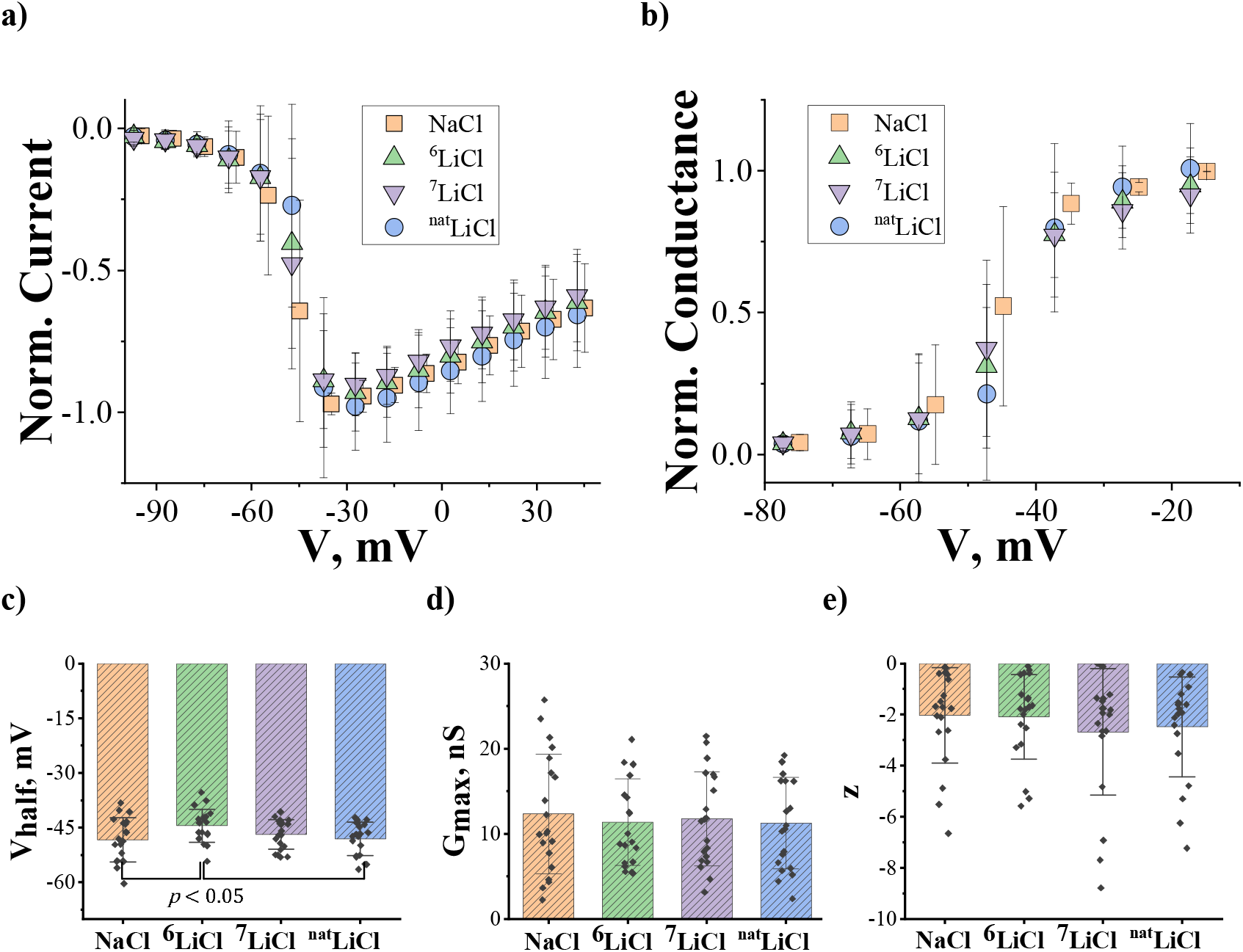
Normalized current/conductance profiles and Na^+^ channel parameters from iPSC-derived cortical neurons upon constant perfusion of NaCl, ^6^LiCl, ^7^LiCl, ^nat^LiCl. **Figure 3. a)** Corresponding *I*−*V* profiles for Na^+^ and Li^+^ ions, where current magnitudes were normalized to the maximum inward current measured solely under Na^+^ perfiision. **b)** Corresponding *G* − *V* profiles for Na^-^ and Li^+^ ions, where conductance magnitudes were normalized to the maximum conductance measured solely under Na^+^ perfiision. **c)** *V*_half_ − the half-maximum activation potential, cl) G_max_ − the maximal conductance, **e**, *z* − apparent valence of the gating charge. The symbols in panels **a)** and **b)** and columns in panels **c)-e)** represent the mean, the data point (black dots) in panels **c)-e)** represent individual measurements, and bars represent standard deviation (SD). Statistical significance for individual cells was evaluated using one-way repeated measures ANOVA. The number of replicates (individual cells successfully patched) is *n* = 20.

### Comparison of Na^+^ channels from SH-SY5Y and iPSC-derived cortical neurons

Having obtained results for iPSC-derived cortical neurons and SH-SY5Y, we proceeded to compare these cell lines with each other to see how different the Na^+^ channel parameters are.

The fitted parameters (*V*_half_, *G*_max_, *z*) of the cunent-voltage curves for SH-SY5Y cells were compared to parameters for iPSC-derived cortical neurons [Figure **4a)-4c**)] within the same buffer. Differences in these parameters indicate differences in isoforms and quantity of Na^+^ channel expressions which is expected to be found between two distinct cell lines. The half-activation potential is more negative in iPSC cells (see Table 1) in every buffer (except ^6^LiCl) [Figure **4a)**, peach, violet, and blue]. The maximum conductance was found to be much larger in iPSC-derived cortical neurons as visible in Figure **4b)** (with a significance level of *p* < 0.001 for NaCl, ^6^LiCl and ^7^LiCl and with significance level of *p* < 0.01 for ^nat^LiCl). The apparent valence of the gating charge was found to be significantly different in all buffers with the level of *p* < 0.001 [Figure **4c**)].

**Fig. 4:**
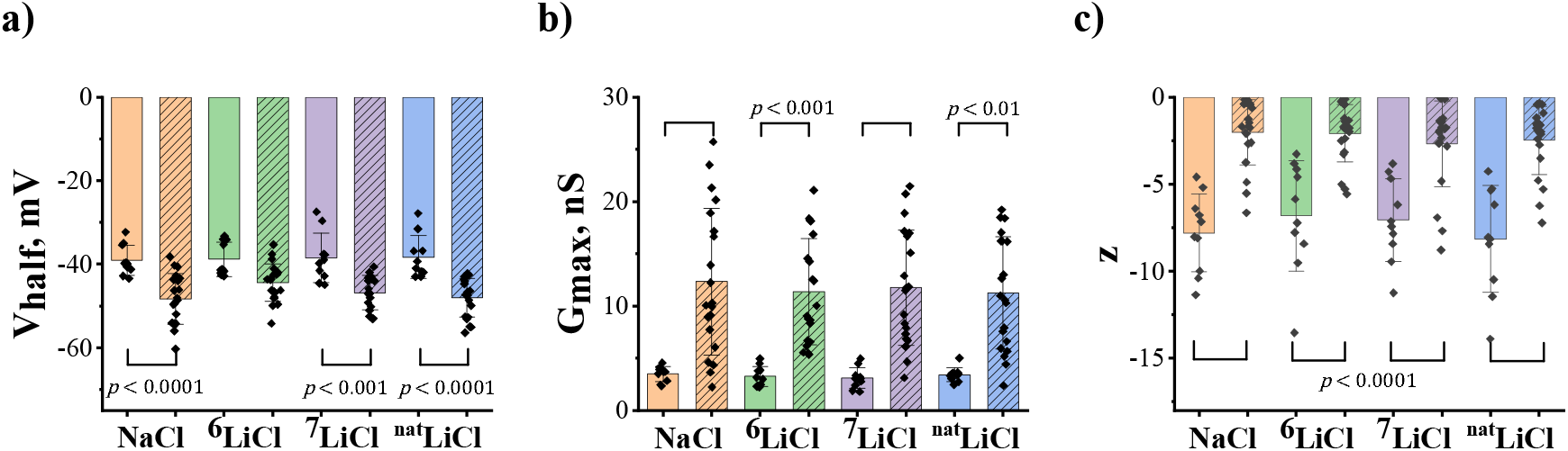
Na^+^ channels parameters from SH-SY5Y and iPSC-derived cortical neurons in constant perfusion of NaCl, ^6^LiCl, ^7^LiCl, ^nat^LiCl. **Figure 4. a)** *V*_half_ - the half-maximal activation potential, **b)** *G*_max_ - the maximal conductance, **c)** *z* - the apparent valence of the gating charge. Statistical significance was evaluated by one-way ANOVA with Bonferrom’s post hoc test (solid columns represent SHSY-5Y cells, dashed columns represent iPSC-deiived cortical neurons; points represent the individual measurements, colored boxes represent the mean, and bars represent SD). The number of replicates (individual cells successfully patched) is *n* = 20 for iPSC-derived cortical neurons and n = 10 for SH-SY5Y cells.

**Table 1.** The half-maximal activation potential for SHSY-5Y cells and iPSC-derived neurons in constant perfusion of 135 mM of NaCl, ^6^LiCl, ^7^LiCl, ^nat^LiCl.

|  |  | NaCl | <sup>6</sup>LiCl | <sup>7</sup>LiCl | <sup>nat</sup>LiCl |
| --- | --- | --- | --- | --- | --- |
| $V_{\text{half}} \pm \text{SD}, \text{ mV}$ | iPSC-derived cortical neurons | $-48.3 \pm 6.0$ | $-44.4 \pm 4.5$ | $-46.9 \pm 4.1$ | $-48.1 \pm 4.6$ |
| | SHSY-5Y cells | $-39.0 \pm 3.6$ | $-38.9 \pm 4.1$ | $-38.5 \pm 5.9$ | $-38.3 \pm 5.1$ |

## Discussion

Several previous reports exploring the differences between Li^+^ isotopes in various experimental settings, including animal and cellular studies^4,5,11,13^, vary in their conclusions and emphasize the importance of further studies to help identify the molecular target for Li^+^ isotope differentiation. Considering the most striking large and opposite effects of Li^+^ isotopes observed recently in the electrical response in animal brain tissues^4^, we hypothesize that fast electrochemical processes could be responsible for such differences. Therefore, ion channels involved in neuronal signal propagation and transmission are of interest in this regard, including Ca^2+^, Na^+^ and K^+^ channels.

Li^+^ is well-known to pass through Ca^2+^ channels^26^ and Na^+^ channels^14,16^ while possibly inhibiting current in K^+^ channels^27^. In this work, we studied Na^+^ channels as a potential target for Li^+^ isotope differentiation. Na^+^ channels are known to be one of the primary pathways for Li^+^ entry into cells^14–18^, but the particular isoform of the channel as well as the details of the mechanism of how Li^+^ affects the Na^+^ channel remain unknown. Studies on bacterial Na^+^ channels have shown that Na^+^ to Li^+^ substitution leads to a decrease in current through the Na^+^ channels^17,18^ while research on rat skeletal muscle showed no difference between Na^+^ and Li^+ 15^. These findings align with known differences in the selectivity filters of different channels^28^.

In this work, we used two cell lines: SH-SY5Y and iPSCs-derived cortical neurons. The SH-SY5Y cell line has been well-studied and characterized^22^. In one study, it was found that these cells express a high level of different Na^+^ channel isoforms: Nav1.2, Nav1.3, Nav1.4, and Nav1.5, with Nav1.7^29^ as the most highly expressed. Another study found that SH-SY5Y cells express Nav1.2, Nav1.3 and Nav1.9^30^, with Nav1.7^30^ being the most highly expressed. We, therefore, assume that our patch-clamp recordings in SH-SY5Y primarily reflect Nav1.7 channels, as indicated in Blum *et al*. 2002^30^ and Vetter *et al*. 2012^29^. In the iPSCs-derived cortical neurons, Nav1.1 is predominant in inhibitory gamma aminobutyric acid-ergic (GABAergic) interneurons^31,32^, Nav1.2 prevails in excitatory neurons^33^ and glutamatergic neurons^34^ and Nav1.6 is expressed in neocortical excitatory neurons^35^. In our work, we followed previously published protocols^36,37^ where iPSC-derived cortical neurons were grown and characterized, producing primarily excitatory and glutamatergic neurons. Approximately 80% of the neurons in our culture are expected to be glutamatergic, making Nav1.2 the most likely isoform recorded during the patch-clamp experiments.

We studied Li^+^ ions passage through Na^+^ channels in the two aforementioned cellular models. Instead of selecting physiologically relevant Li^+^ concentrations, and in line with concentrations considered in previous works^17,18^, we opted to consider an acute setting by fully substituting Na^+^ with Li^+^ in order to magnify any differences that might exist. We found that the two Li^+^ isotopes pass through the Na^+^ channels in both neuronal models while Cs^+^ and K^+^ ions do not permeate through these channels, as would have been expected based on previous studies on other isoforms of Na^+^ channels and bacterial Na^+^ channels^15,17,18^. We found no differences between Na^+^ and any Li^+^ (isotope or natural) in any of the parameters (half-activation potential, maximum conductance and apparent valence of the gating charge) describing the *I-V* characteristics in the mammalian SH-SY5Y model cell line. This result indicates that mammalian Na^+^ channels do not show selectivity between Na^+^ and Li^+^ in contrast with bacterial Na^+^ channels^17,18^ where Li^+^ (natural isotopic abundance) has a lower permittivity than Na^+^ through Na^+^ channels. Our results are in agreement with another study carried out on mammalian Na^+^ channels^15^ in HEK293 cells where it was found that the Li^+^ and Na^+^ ions display no difference in their permittivity through the Na^+^ channels. Na^+^ channels in iPSC-derived cortical neurons showed differences in the half-maximum activation potential between Na^+^ and ^6^Li^+^ and between Li^+^ and ^6^Li^+^ ions. We saw no drastic differences as those found in bacterial channels, where the difference between Na^+^ and Li^+^ induced currents is nearly 2-fold^17,18^. The different results between the two cellular models must thus be attributed to the different Na^+^ channels subunit isoforms that have distinct selectivity filters^28^. On one hand, since Na^+^ channels are expressed ubiquitously, even relatively small changes in some parameters could impact various behavioural or therapeutic outcomes, their activity can be different in different cell lines as we report here for SH-SY5Y and iPSC cells. On the other hand, there is no differentiation between Na^+^ and Li^+^ or Li^+^ isotopes in SH-SY5Y cells and very small differentiation in iPSC-derived cortical neurons used in the study which cannot explain the large and opposite effects observed in the previous tissue^4^ or behavioural^3^ studies. Thus, our findings, within the methodology and experimental conditions used in this work, suggest that Li^+^ regulation through Na^+^ channels would likely not be the primary molecular mechanism responsible for Li^+^ isotope differentiation observed in other studies^3-11^. Moreover, our findings that Na^+^ channels in mammalian cells are not selective in ion passage compared to those reported in bacteria^17,18^ suggest an intriguing research direction which we have not explored in the present work, but would constitute a natural extension, namely: might bacterial channels differentiate between Li isotopes? What properties of bacterial channels would then govern the observed differences? Beyond the interest in Li isotope effects that may exist in neuroscience, these questions motivate new avenues for further investigations into the detailed molecular mechanisms underlying ion channel selectivity.

Our data provide valuable insights into the permeability of Na^+^ channels in comparison to Li^+^ ions and its isotopes. However, several limitations must be acknowledged to contextualize the findings and help guide future research. Our study’s primary goal was to determine whether Na^+^ channels alone can explain the large differences observed in Li^+^ isotope effects on field excitatory postsynaptic potentials (fEPSPs)^4^. To be focused on this goal we constrained our experimental design. First, the experimental conditions involved the full substitution of Na^+^ with Li^+^ and conducting experiments at room temperature (RT). These conditions, while useful for isolating effects, do not reflect physiological environments. At RT, channel and enzyme rate constants are reduced compared to physiological temperatures, potentially altering ion channel dynamics and kinase interactions. Future studies at physiological temperatures could provide more accurate insights. Second, our findings are based on two neuronal models: SH-SY5Y neuroblastoma cells and iPSC-derived cortical neurons. While these models provide complementary perspectives, they do not fully represent the diversity of Na^+^ channel isoforms in human tissues, some of which may differ significantly from those present in the brain. Furthermore, the study investigated all Na^+^ channels present in these cells without targeting specific isoforms. While this broad approach provides general insights, it limits specificity regarding individual channel isoforms and their unique roles. Additionally, the iPSC-derived neurons exhibit diverse and complex morphologies, which can influence electrophysiological recordings and ion channel behavior. Na^+^ channels were assumed to be the primary conductors of Li^+^ based on established literature^14–18^. However, this assumption may oversimplify the complexity of Li^+^ transport and interactions in cellular environments and hide some of the Li^+^ effects. Lastly, to preserve cell integrity during seven buffer exchanges, we limited current recordings to high positive voltages. Notwithstanding these limitations, we believe that our study successfully addresses its primary question: whether Na^+^ channels can explain the large differences observed in Li isotope effects on fEPSPs^4^.

In summary, we used whole-cell patch-clamp techniques to study the effect of Li^+^ and Li^+^ isotopes on the electrical activity of Na^+^ channels in SH-SY5Y neuroblastoma cells and human iPSC-derived cortical neurons. We found no statistically significant differences between Na^+^ and Li^+^ or Li^+^ isotopes in SH-SY5Y cells. On the other hand, we detected small differences in the half-maximum activation potential between Na^+^ and ^6^Li^+^ and between Li^+^ and ^6^Li^+^ in iPSC-derived cortical neurons. Our comparison study revealed significant differences in half-activation potential, maximum conductance and apparent valence of gating charge between SH-SY5Y and iPSC-derived cortical neurons for Na^+^ and Li^+^ ions, supporting the idea that these cell lines have different abundance of different Na^+^ channel isoforms, emphasizing the importance of considering the optimal cell model for a specific research study.

## Materials and Methods

### SH-SY5Y and iPSC-derived cortical neurons maintenance

SH-SY5Y cells were cultured as previously described^29^ with some modifications described below. Briefly, the cells were maintained at 37°C in Dulbecco’s Modified Eagle’s Medium supplemented with 10% fetal bovine serum, 100 μg/ml penicillin/streptomycin mix. Approximately 2 days after seeding, cells were plated at low density on Poly-l-Lysine coated glass coverslips, contained in a 12-well plate, allowed to grow for at least 2 days and up to 10 days and then used for electrophysiological recordings. Media changes were performed every 3-4 days. SH-SY5Y cells expressing endogenous Na^+^ channels channel typically exhibited a maximum peak Na^+^ current of ∼0.4 nA, which is in alignment with previous studies^29,38,39^. The measured membrane capacitance range for SHSY-5Y cells was 5-10 pF.

iPSC-derived neural stem cells (NSCs) were purchased from Alstem (hNSC11). NSCs were cultured and differentiated into cortical neurons according to the following protocol^36,37^. NSCs were thawed in neural progenitor media (NPM) (StemCell Technologies, 05833) with Supplement A, Supplement B added, and 10 μM ROCK inhibitor y-27632 (Sigma-Aldrich), and then seeded onto Matrigel (Corning) coated culture vessels. After 24 hours, the media was replaced to remove the ROCK inhibitor. 48 hours prior to passaging the NSCs for differentiation, the NPM was exchanged to remove supplement B to prevent rapid overgrowth of NSCs during neuron differentiation. For passaging, the NSCs were detached using Accutase (ThermoFisher, A11105-01) and seeded onto glass coverslips, coated with Poly-L Ornithine (Sigma-Aldrich, P4957) and laminin (ThermoFisher, 23017015), contained in 12-well plate. Cells were seeded at a density of 50,000 to 75,000 NSCs per well in NPM containing only Supplement A. Then, 24 hours after seeding, neural differentiation media (BrainPhys Media (StemCell Technologies, 05790), SM1 (StemCell Technologies, 05711), N2A (StemCell Technologies, 07152), 20 ng/mL BDNF (StemCell Technologies, 78005), 20 ng/mL GDNF (StemCell Technologies, 78058), 200 nM L-Ascorbic acid (StemCell Technologies, 100-1040), 1 mM cAMP (StemCell Technologies, 73886) was added to each well. Half media changes were performed every 2-3 days. After 6-8 weeks of neuronal differentiation, cells were ready for patch-clamp electrophysiology. Cortical neurons expressing endogenous Na channels channel typically exhibited a maximum peak Na^+^ current of ;1 nA. The measured membrane capacitance range for iPSC-derived neurons was 15-30 pF.

### Electrophysiology

Patch clamp pipettes were fabricated on a PC-10 two-stage electrode puller (Narishige USA, Inc., Glen Cove, NY) using Sutter borosilicate glass with Filament (Su-BF150-86-10). The pipette resistance was measured to be 4 to 8 MΩ when filled with standard pipette solution (in mM: CsCl (120), NaCl (5), MgATP (5), HEPES (20), EGTA (10), CaCl_2_ (2), pH was adjusted to 7.3 with CsOH). Whole-cell voltage-clamp recording was performed at room temperature (∼;22°C) using an Axopatch 200B (Axon Instruments, Union City, CA) and Clampex software (sampling rate 250.00 kHz). Some restrictions were applied to take into account differences between individual cells, signal-to-noise ratio, and quality of the signal. Cells were not accepted for recording if the initial seal resistance was less than 2 GΩ or if the peak Na^+^ current was less than 0.2 nA for SH-SY5Y cells and less than 0.5 nA for iPSC cells. Additionally, series resistance was measured throughout each experiment and the recording was only accepted if the series resistance varied by less than 20%. Current parameters were chosen according to the most large and frequent but stable cases. Compensation of the capacitance transients and leak subtraction was performed using a programmed P/4 protocol. Membrane currents were filtered at 5 kHz.

All cells were voltage clamped in the whole cell configuration. Current-voltage data were collected by recording responses to a consecutive series of step pulses from a holding potential of − 100 mV at intervals of +10 mV beginning at − 100 mV and up to +50 mV (with pulse duration of 10 ms and pulse frequency of 1 Hz). Data collection was initiated approximately 5 min after break-in when control Na^+^ currents had stabilized after intracellular perfusion with pipette solution. Data was always recorded during continuous perfusion of the clamped cell with the extracellular solution. Permeability to different cations was tested by recording a current-voltage (*I* – *V*) sequence, first in the control Na^+^ solution, and then during perfusion with a sequence of test cations solutions, and again after replacement with the control Na^+^ solution. Na^+^ currents often progressively decreased over time (termed “rundown”) using this method. This effect was estimated by fitting the decay of current during the control perfusions to a linear equation that was interpolated over the time course of the experiment.

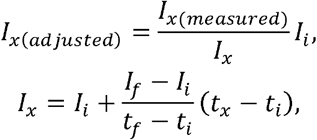

where *I*_*x*_ is the interpolated current, *t*_*x*_ is the time when the cunent of interest was recorded, *I*_*i*_ is the is the cunent amplitude measured during the first control NaCl perfusion (initial), *I*_*f*_ is the cunent amplitude measured during the last control NaCl perfusion (final), *t*_*i*_ is the is the time of the first control NaCl perfusion (initial), *t*_*f*_ is the time of the last control NaCl perfusion (final). Using this estimation, the rundown was compensated.

### Solutions and Chemicals

Li salts, ^6^LiCl (95% ^6^Li) and ^7^LiCl (99% ^7^Li), as well as ^nat^LiCl salt (with natural abundance – ^6^Li at 7.49% and ^7^Li at 92.51%), were purchased from Sigma-Aldrich.

The intracellular (pipette) solution contained (in mM) CsCl (120), NaCl (5), MgATP (5), HEPES (20), EGTA (10), CaCl_2_ (2) and pH was adjusted to 7.3 with CsOH. Osmolarity was measured to be ∼270 - 280 mOsm. The intracellular solution was prepared as a stock, without adding MgATP and frozen. MgATP was added just before the recording. The internal solution was kept on ice for the entire duration of the experiment, to prevent MgATP degradation. The external solutions contained (in mM) either NaCl, LiCl (natural or isotopes), CsCl or KCl (135), HEPES (10), CaCl_2_ (2), KCl (5), MgCl_2_ (1), Glucose (10) and the pH was adjusted to 7.4 with KOH. Osmolarity was adjusted with glucose to be ∼290 - 300 mOsm. The external solution was prepared as 10X without adjustment for osmolarity and pH and frozen for storage. pH and osmolarity were adjusted just before the experiment. All solutions were filtered with a 0.22-μm filter before use. The external recording solution in the experimental chamber was continuously exchanged by a gravity-driven arrangement at the rate of ∼1 ml/min.

## Data analysis

Data were analyzed by using the Easy Electrophysiology and Origin software. Current-voltage curves were fitted using the Ohm’s law/Boltzmann expression^40^:

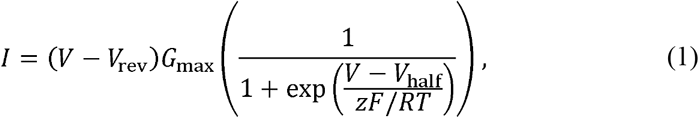

where *I* is the current amplitude for the applied potential *V, V*_rev_ is the reversal potential, *G*_max_ is the maximal conductance, *z* is the apparent valence of the gating charge, *V*_half_ is the half-maximal activation potential, *R, T* and *F* are the universal gas constant, absolute temperature, RT and the Faraday constant, respectively, 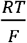 is 25.4 mV (room temperature was assumed to be 295 K). Since the bathing solution was changed during the experiments, the correction for liquid junction potential was conducted after the experiment and for each cation condition (−; 0.2 to 7.3 mV). Liquid junction potential was calculated according to the stationary Nernst-Planck equation^41^ using LJPcalc (RRID:SCR_025044).

Due to experimental design, we did not measure *V*_rev_. However, we estimated it using Ohm’s law/Boltzmann expression. We report these data in Supplementary Material, Figure S3. Upon analysis, we found that in some cases, *V*_rev_for iPSCs-derived cortical neurons was overestimated in the positive direction (being greater than 250 mV in some cases). We attribute this to several well-known space clamp properties in iPSCs-derived cortical neurons, including the presence of ion channels in both the axon and cell body^42^, discrepancies between the command voltage and the actual voltage across the membrane in case of large current flow^43^, and difficulty in uniformly controlling voltage in cells with complex structures^44^. We believe these issues with *V*_rev_ stem from the factors mentioned and not from experimental quality, as data from SH-SY5Y cells remained highly consistent.

Statistical significance for individual cells was evaluated using One-Way Repeated Measures ANOVA. Statistical significance for SH-SY5Y and iPSCs-derived neurons comparison was evaluated by one-way ANOVA with Bonferroni’s post hoc test, with data reported as Mean ± SD. The number of replicates (individual cells successfully patched) is n =10 − 12.

## Acknowledgments

I.B. acknowledges the Waterloo Institute for a Nanotechnology (WIN) Nanofellowship. E.V.P. acknowledges the Waterloo Institute for a Nanotechnology (WIN) travel award. We appreciate stimulating and useful discussions with all members of the Waterloo Quantum Biology team. We appreciate Dr. N. Murugan and Dr. N. Rouleau’s lab for generous permission to use their equipment for pipette preparation.

## Author Contributions

Conceptualization, E.V.P., I.B., M.J.P.G. and Z.L.; methodology, E.V.P., K.S., J.D.L. and M.A.B.; validation, E.V.P.; formal analysis, I.B., M.J.P.G.; investigation, I.B., J.D.L; resources, M.A.B.; data curation, I.B.; writing—original draft preparation, I.B.; writing—review and editing, I.B., M.J.P.G., K.S., J.D.L., E.V.P., M.A.B., Z.L.; visualization, I.B.; supervision, Z.L., E.V.P. and M.J.P.G.; project administration, Z.L.; funding acquisition, Z.L. and M.J.P.G.. All authors have read and agreed to the published version of the manuscript.

## Data Availability Statement

The datasets generated during and/or analyzed during the current study are available from the corresponding author upon reasonable request.

## Funding

This research was funded by New Frontiers in Research Fund (NFRF), grant number 50383-10011 (Z.L., M.J.P.G.); Natural Sciences and Engineering Research Council of Canada (NSERC) grant number 50503-11497 (Z.L.); National Institute of General Medical Sciences (NIGMS), grant number R35GM139615 (E.V.P.).

## Conflicts of Interest

The authors declare no conflicts of interest.

## Notes

### Competing Interest Statement

The authors have declared no competing interest.

