## Supplementary materials for "Effects of Natural Lithium and Lithium Isotopes on Voltage Gated Sodium Channel Activity in SH-SY5Y and IPSC Derived Cortical Neurons"

**a)**

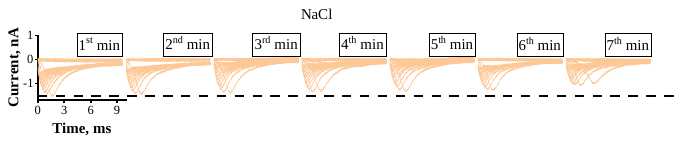

**b)**

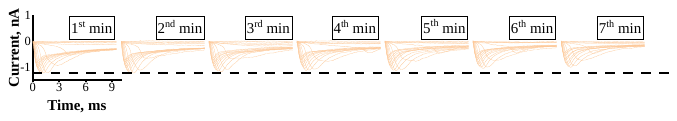

**c)**

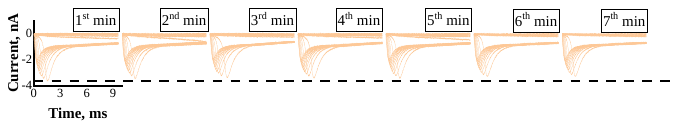

**Figure S1. Rundown variability.** Here we present three examples of control rundown experiments – each cell was perfused with Na^+^ solution for 7 minutes. Given the variability between the cells, we decided to apply linear interpolation and compensate for the rundown for each cell and not on average.

| **a)** | **b)** |
| --- | --- |
| 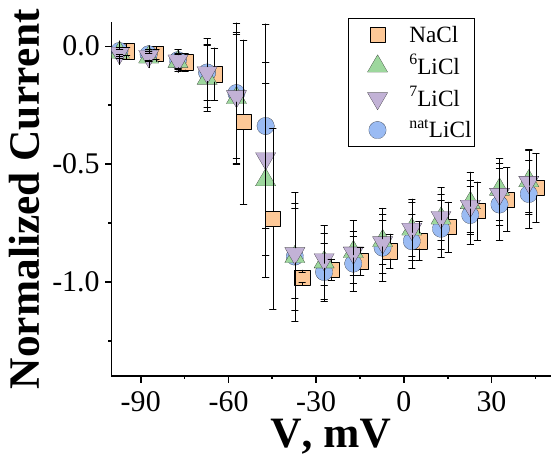 | 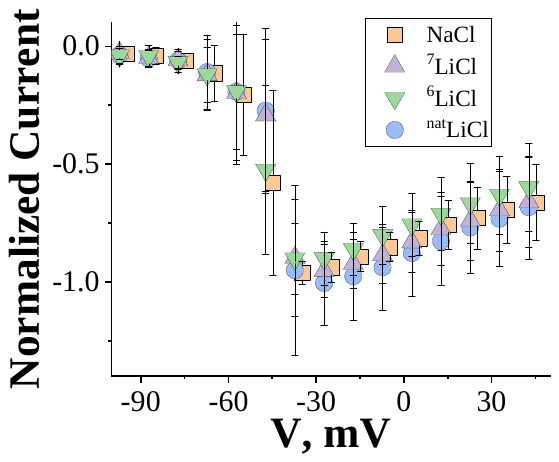 |
| **c)** | **d)** |
| 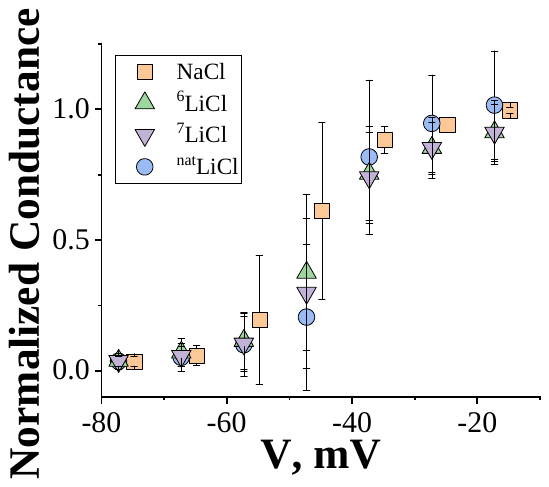 | 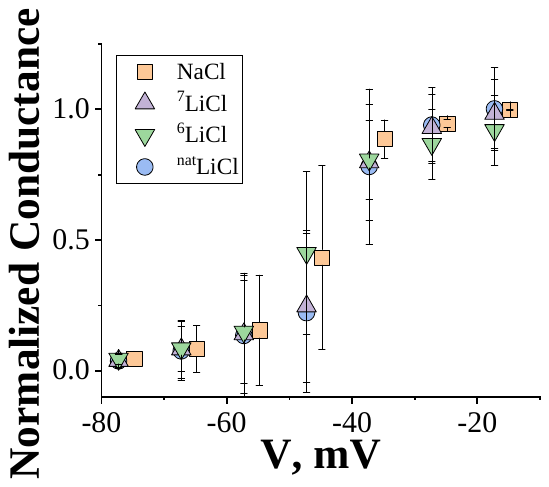 |

**Figure S2. Randomized order of ^6^Li and ^7^Li.** Here we present 2 groups of the experiments – with ^6^LiCl preceding ^7^LiCl [a) and c)] and with ^7^LiCl preceding ^6^LiCl [b) and d)]. These groups were not significantly different in terms of $I-V$, $G-V$ and parameters extracted from the Boltzmann fit (not shown). Therefore, we decided to combine these two groups for the main text.

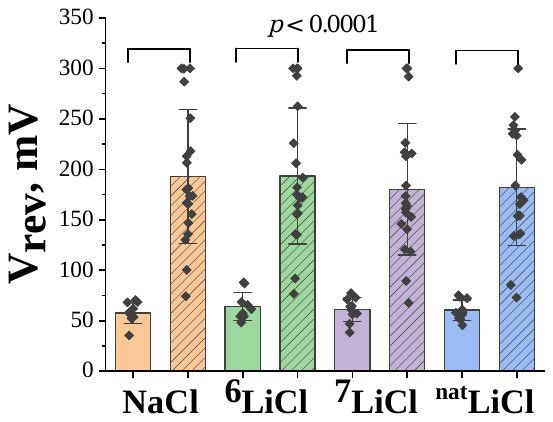

**Figure S3.** $\boldsymbol{V}_{\mathbf{rev}}$ **approximation** **for SH-SY5Y cells and iPSC-derived cortical neurons.** Statistical significance was evaluated by one-way ANOVA with Bonferroni’s post hoc test (solid columns represent SHSY-5Y cells, dashed columns represent iPSC-derived neurons; points represent the individual measurements, columns represent the mean, bars represent SD).
